# Unraveling the genetic legacy of commercial whaling in bowhead whales and narwhals

**DOI:** 10.1101/2024.01.04.574243

**Authors:** Evelien de Greef, Claudio Müller, Matt J. Thorstensen, Steven H. Ferguson, Cortney A. Watt, Marianne Marcoux, Stephen D. Petersen, Colin J. Garroway

**Author notes:** Corresponding author: Evelien de Greef.

## Abstract

Commercial whaling decimated many whale populations over several centuries. Bowhead whales (*Balaena mysticetus*) and narwhal (*Monodon monoceros*) have similar habitat requirements and are often seen together in the Canadian Arctic. Although their ranges overlap extensively, bowhead whales experienced significantly greater whaling pressure than narwhals. The different harvest histories but similar habitat requirements of these two species provide an opportunity to examine the demographic and genetic consequences of commercial whaling. We whole-genome resequenced Canadian Arctic bowhead whales and narwhals to delineate population structure and reconstruct demographic history. Bowhead whale effective population size sharply declined contemporaneously with the intense commercial whaling period. Narwhals instead exhibited recent growth in effective population size, reflecting limited opportunistic commercial harvest. Although the genetic diversity of bowhead whales and narwhals was similar, bowhead whales had more genetic diversity prior to commercial whaling and will likely continue to experience significant genetic drift in the future. In contrast, narwhals appear to have had long-term low genetic diversity and may not be at imminent risk of the consequences of the erosion of genetic diversity. This work highlights the importance of considering population trajectories in addition to genetic diversity when assessing the genetics of populations for conservation and management purposes.

## 1. Background

Intense commercial whaling in recent centuries brought many whale populations to the brink of extinction [1], potentially leading to long-lasting negative effects on their genetic diversity through genetic drift. This is concerning because genetic diversity underlies population resilience and the capacity to adapt to future environmental change. Addressing the consequences of whaling on genetic diversity is important for understanding the genetic health of populations, yet it remains largely unaddressed in many species (but see [2,3]). Although some whale populations started to recover in number of individuals after whaling moratoriums were implemented (e.g., Canada’s ban on commercial whaling of bowhead whales in 1915 [4], a global whaling moratorium in 1982 [5]), genetic diversity can take millennia to recover because of the slow rate at which mutations accumulate [6]. Continued loss of genetic diversity can eventually put a population at risk for inbreeding depression, consequently affecting fitness and limiting population growth (e.g., [7]). Understanding evolutionary processes in whale species that faced commercial harvest pressure provides context for making conservation decisions for vulnerable populations.

Although the cessation of commercial whaling protects populations from further industrial harvest, the continued recovery and future status of whale populations are threatened by climate change [8–10]. The Arctic is warming four times faster than the global average [11], highlighting a need to understand present structure and past population changes in Arctic whales. Narwhals (*Monodon monoceros*) and bowhead whales (*Balaena mysticetus*) are endemic to the Arctic and are integral to Arctic ecosystems as both predators and prey of other species. Due to their similar habitat requirements, both species are often seen together. Changes in their abundance, distribution, and genetic health may greatly impact marine wildlife and local communities. The projected loss of sea ice due to climate change will affect populations through habitat loss [12,13], changes in prey distribution [14,15], increased predation pressure from killer whales (*Orcinus orca*) [16], and increased ship traffic [17]. As ice-adapted animals, narwhals and bowhead whales will need to adjust quickly to changing environments, which may be limited by potential genetic consequences carried over from commercial whaling.

While both narwhal and bowhead whale ranges overlap in the Canadian Arctic, they have very different current and past population sizes and have faced different intensities and lengths of commercial whaling. Narwhals were generally opportunistically harvested [18] as they were considered a supplementary source of oil and for their ivory tusks [19], and because they were more difficult to hunt than other whale species [20]. Commercial narwhal harvests in the eastern Canadian Arctic were infrequently mentioned in whaling records until the 19^th^ century, followed by a reported 558 to 754 narwhals harvested from the Davis Strait in the 19^th^ to early 20^th^ century [20]. By contrast, bowhead whales were popular targets for whalers [21] due to their value from high oil content, baleen, and large size. Bowhead whales were overharvested in succession from the eastern North Atlantic regions (Svalbard-East Greenland), the areas within eastern Canadian Arctic (Eastern Canada-West Greenland), the Pacific Ocean (Bering-Chukchi-Beaufort), and lastly the Okhotsk Sea population. In eastern Canada and west Greenland, it is estimated that over 55,900 bowhead whales were commercially harvested between the 16^th^ to 20^th^ centuries [22]. While bowhead whales in the eastern Canadian Arctic have made recoveries in numbers [23], the effects of genetic drift may still cause declines in genetic diversity. As long-lived animals (lifespan reaching over 100 years for narwhal [24]; and over 200 years for bowhead whale [25]) with long generation times, these whale populations may take a long time to recover from declines in genetic diversity, which is especially concerning given their need to respond relatively quickly to changing climates.

Reconstructing demographic history can help explain current levels of genetic diversity by evaluating the severity and timescale of population bottlenecks and expansions [26]. In the context of the history of whaling, demographic models can be used to examine changes in effective population size (*N_e_*) coinciding with intense harvest (e.g., [2,3]). The *N*_e_ of a population is an estimate of the strength of genetic drift and the efficiency of selection [27]. Although *N*_e_ is related to the number of individuals in a population, *N*_e_ is usually a smaller value than total population abundance because not every individual contributes to the population equally. Contemporary estimates of *N*_e_ are important for quantifying the magnitude of genetic drift and genetic diversity [27,28] and can be an indicator of genetic risk.

To investigate genomic impacts of whaling and improve our understanding of population genomics in Arctic whales, we analyzed whole-genome data from narwhals and bowhead whales sampled across the eastern Canadian Arctic. First, we examined population structure to identify genetic clusters and estimated various metrics of genetic diversity. We then used multiple methods to reconstruct demographic history from contemporary years encompassing industrial whaling into the Pleistocene era to examine how population declines or expansions may have been affected by large-scale events.

## 2. Materials and Methods

### (a) Resequencing data

Tissue samples were obtained from harvested narwhals (*n* = 62) and bowhead whales (*n* = 21) in the eastern Canadian Arctic by Inuit subsistence hunters between 1982 and 2020 (locations shown in Figure 1a, Figure 2a, and listed in Table S1). After DNA extraction with the Qiagen DNeasy Blood & Tissue kit, samples were whole-genome sequenced with Illumina NovaSeq. We trimmed raw sequences with *Trimmomatic* v0.36 [29], then aligned the reads to their respective reference genomes (narwhal genome from NCBI Accession GCA_005190385.2, [30]; bowhead whale genome from www.bowhead-whale.org, [25]) using *BWA* v0.7.17 [31]. Through *Picard* v2.20.6 [32], we removed duplicate reads and added read group information. Additionally, we filtered reads to include only primary alignments mapped in proper pairs through *SAMtools* v1.9 [33]. To avoid downstream biases in variant calling due to sample coverage variation (3 – 24x for narwhal; 7 – 19x for bowhead whale), we used *GATK* v4.1.2 [34] to down-sample select samples to the modal coverage (10x for narwhal, Table S2; 11x for bowhead whale, Table S3).

**Figure 1.**
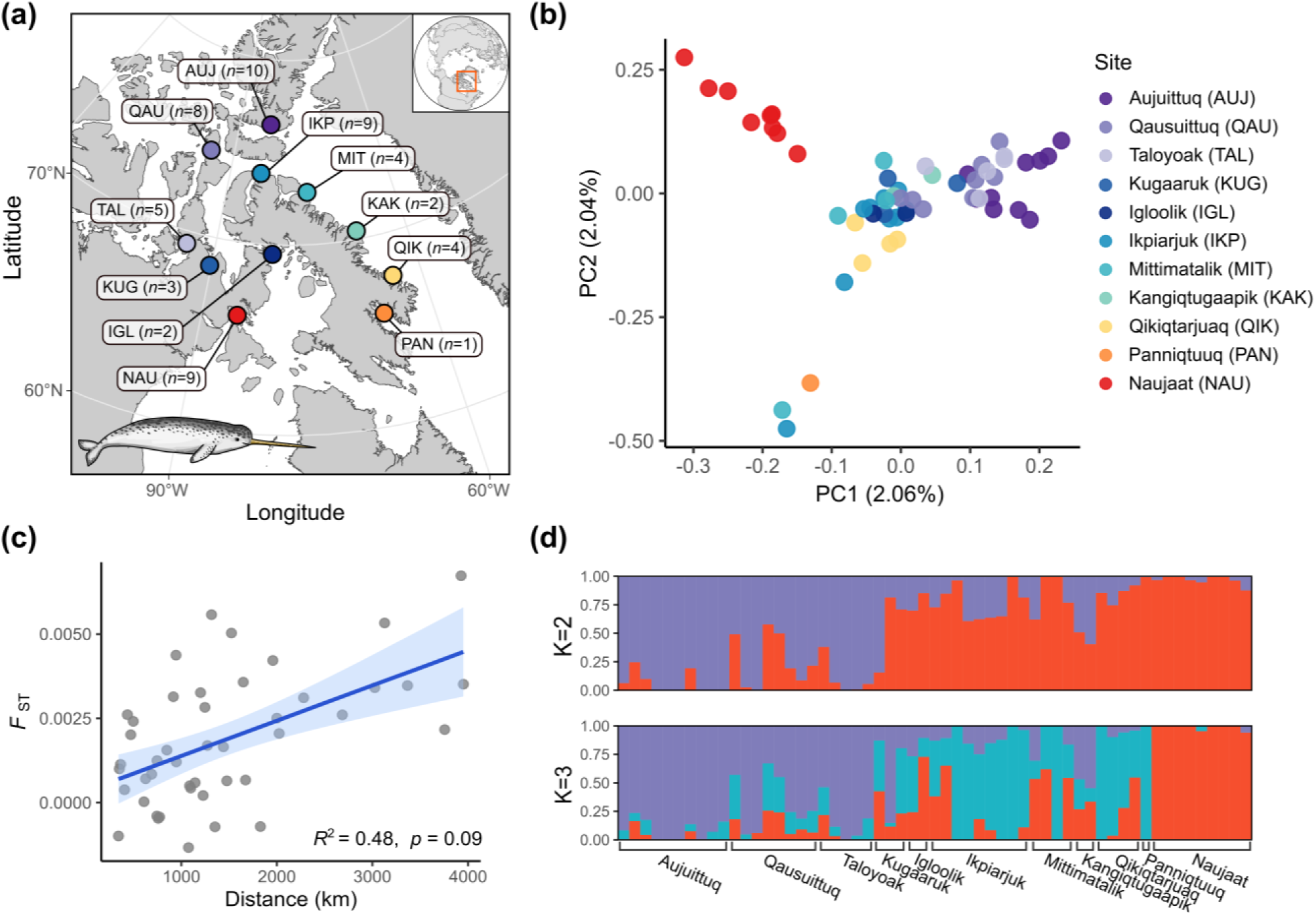
(a) Map of sites with number of eastern Canadian Arctic narwhal samples next to site ID. Population substructure in narwhal (*n* = 57) is shown by (b) regional clustering in PCA and (c) isolation-by-distance, with a positive correlation between pairwise *F*_ST_ and distance between sites, and (d) admixture models with K = 2 and K = 3 ancestral groups.

**Figure 2.**
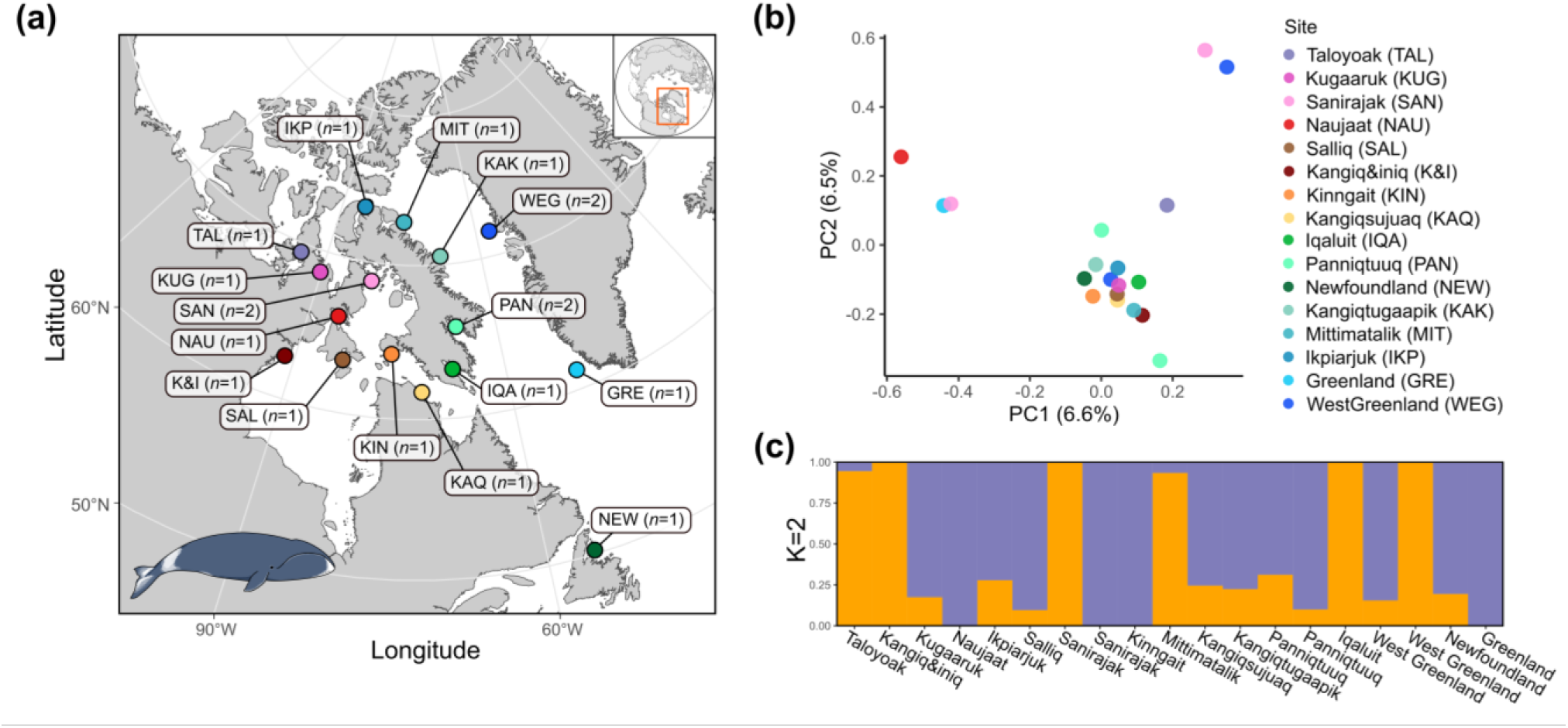
(a) Map of sites with number of Canadian Arctic bowhead whale samples next to site ID. Lack of clear population structure in bowhead whale (*n* = 19) shown by (b) PCA and (c) ancestral admixture with K = 2 ancestral groups (individuals are ordered by longitude).

Genomic variants were called using *Platypus* v0.8.1 [35], then filtered to high-quality autosomal datasets of single-nucleotide polymorphisms (SNPs), removing sites of QUAL < 50, MQ < 40, QD < 4, missingness > 0.25, non-biallelic sites, small scaffolds (< 100 kb), and sex-linked sites with *VCFtools* v0.1.17 [36]. See supplemental for details on identifying sex-linked scaffolds. Using *PLINK* v1.9 [37], we identified kin pairs (pi-hat ≥ 0.25, the threshold for 2^nd^ degree relatives) and removed the individual with more missing data from each pair. We also removed any duplicate samples, and samples that had more than 30% overall missingness (Table S2, Table S3).

### (b) Population structure

SNPs in population structure analyses were further filtered to remove loci under possible selection and non-random association of loci (Hardy-Weinberg equilibrium heterozygous frequency threshold > 0.6, minor allele frequency < 0.05, and linkage disequilibrium *r*^2^ > 0.8; Table S4). We examined population structure using Principal Component Analysis (PCA) with R-package *pcadapt* v4.3.3 [38] and ancestral admixture through the snmf function in R-package *LEA* v3.3.2 [39]. We analyzed results with R version 4.2.1 [40].

In narwhal specifically, we assessed patterns of isolation-by-distance using sites with at least two samples. Here, we estimated pairwise genetic differentiation (*F*_ST_) following Reich et al. [41]’s method to reduce biases from small sample sizes, and measured distances over water between sites with R-package *marmap* v1.0.6 [42]. Given the narwhal species’ range does not typically connect between Naujaat and Igloolik, we measured distances between sites accounting for this potential separation. We used a mantel test in R-package *ade4* v1.7.19 [43] to measure the correlation between *F*_ST_ and distance between sites.

### (c) Measures of genetic diversity

A higher the proportion of heterozygous sites indicates greater genetic variability, and thereby higher genetic diversity [44]. We used the same SNP dataset from the population structure analyses and calculated the proportion of heterozygosity for each individual by counting the number of homozygous sites identified with *VCFtools* v0.1.17 [36] and subtracting the count from the total number of SNPs. Additionally, we used *hierfstat* v0.5.11 [45] to estimate overall observed heterozygosity for both species.

Runs of homozygosity (ROH) are long tracts of homozygous genotypes from identical haplotypes inherited from both parents and can be used as an indicator of inbreeding [46]. We estimated ROH in the narwhal and bowhead whale using *PLINK* v1.9 [37], following parameters in Foote et al. [47]: minimum segment length of 300 kb, minimum number of 50 SNPs, density of at least one SNP per 50 kb, gap of 1000 kb, up to 3 heterozygote sites per 300 kb window, and allowing up to 10 missing SNP calls (missing data) in a window. Due to the difference in genome assembly between the two species (narwhal N50 scaffold length = 108 Mb; bowhead N50 scaffold length = 0.9 Mb), we used the largest scaffolds with even SNP densities to minimize potential biases from genome fragmentation for species comparison, using the same scaffolds used in GONE analyses below (largest 12 scaffolds in the narwhal and largest 200 scaffolds in the bowhead whale). We then estimated the proportion of ROH (*F*_ROH_) across the selected genome portion, examining results with minimum ROH lengths of 100 kb.

### (d) Effective population size

We examined trends in *N*_e_ within the last 150 generations, inferred from linkage disequilibrium information with the program *GONE* [48]. Here, we filtered autosomal SNPs to exclude sites out of Hardy-Weinberg equilibrium (heterozygous frequency threshold > 0.6), and minor allele frequency below 0.05. To use an even density of SNPs across the scaffolds, we selected the largest 12 scaffolds in the narwhal genome and the largest 200 scaffolds in the bowhead. Following Kardos et al. [7], we used the model parameters: maximum number of 10K SNPs per scaffold over 1K bins, 1K number of generations for linkage data in bins, and 500 repetitions. To examine changes in *N*_e_ across years, we used generation times of 21.9 years in narwhal and 52.3 years in bowhead whale [49].

While genomic demographic history models are informative for estimating past patterns in a population, they are not ideal for calculating contemporary *N*_e_. To estimate contemporary *N*_e_ for each population, we used the ldNe function in R-package *strataG* v2.5.1 [50,51], which uses linkage information. For this analysis, we removed SNPs with missing data and filtered out sites with minor allele frequency < 0.05, followed by randomly thinning SNPs with *PLINK* v1.9 [37] to create datasets with 25,000 loci. To compare contemporary *N*_e_ with census population sizes (*N*) through *N*_e_ /*N* ratios [52], we pulled abundance estimates from published studies for narwhals from the Canadian High Arctic (*N* = 141,908 [53]), northern Hudson Bay (*N* = 19,200 [54]), and for eastern Canada-west Greenland bowhead whales (*N* = 6,446 [55]).

### (e) Historic changes in effective population size

Multiple approaches are available to estimate changes in *N*_e_ over time. Given that different methods provide information for different timelines of *N*_e_ trajectories [56], we implemented models through *PSMC* and *SMC++* to assess demographic histories reaching further into the past through the Pleistocene era. Broadly, *PSMC* uses a single sample in a pairwise sequentially Markovian coalescent model [57]. Estimating demographic history with this method provides consistency for comparison with results from previous studies. While *SMC++* is also a sequentially Markovian coalescent analysis, it combines site frequency spectrum and linkage information from multiple genomes [58], improving accuracy compared to previous SMC methods for inferring *N*_e_. For both *PSMC* and *SMC++* models, we used previously documented mutation rates (narwhal μ = 1.56 x 10^-8^, bowhead whale μ = 2.69 x 10^-8^; [59]) and the same generation times used when plotting *GONE*. Details for *PSMC* and *SMC++* analyses are listed in the supplemental.

## 3. Results

### (a) Resequencing data

Genomic variant calling resulted in 7.9 and 15.5 million loci in narwhal and bowhead whale respectively, which were filtered to 5.0 and 9.8 million high-quality autosomal SNPs. We removed five narwhal and two bowhead whale samples (duplicates, close kin pairs, or high missingness; Table S2, Table S3), resulting in a total of 57 narwhal and 19 bowhead whale samples for analyses. SNP filters and counts for each analysis are listed in Table S4.

### (b) Population structure

The combination of low overall genetic variation and evidence of genetic subgroups indicate that narwhals in the eastern Canadian Arctic exhibit population substructure. PCA and admixture results support three narwhal subgroups: 1) western Baffin Island (Aujuittuq, Qausuittuq, and Taloyoak), 2) eastern Baffin Island (Panniqtuuq, Qikiqtarjuaq, Kangiqtugaapik, Mittimatalik, Ikpiarjuk, Kugaaruk, Igloolik), and 3) northern Hudson Bay (Naujaat) (Figure 1). Identification of these subgroups remained consistent in PCAs that were filtered for different time frames for harvesting months and removal of three outliers (Figure S1). The most variation was observed between individuals from Aujuittuq and Naujaat (seen along PC1), which span the greatest latitudinal difference among the narwhal sites in this study. However, the small proportion of variance (2%) on the first two PCs indicate low overall individual genomic variation. Population substructure was further supported with clear pattern of isolation-by-distance (*R*^2^ = 0.48) and low overall genetic differentiation (*F*_ST_ values up to 0.007; Figure 1c; Table S5).

No clear population structure was observed in eastern Canadian Arctic bowhead whales (Figure 2). There was no consistency with genetic variation and sampling sites, however, there were some outliers that could potentially represent migrants from other populations. Other principal components (PC3 – 6) further showed a lack of clear clustering (Figure S2). This was also supported by admixture results showing intermixed genetic sources across samples (Figure 2c).

### (c) Measures of genetic diversity

Narwhals exhibited a slightly lower overall proportion of observed heterozygosity (*H*_o_) compared to bowhead whales, however, the difference was minimal: narwhal *H*_o_ = 0.282 (Q1 = 0.281, Q3 = 0.284), bowhead whale *H*_o_ = 0.294 (Q1 = 0.291, Q3 = 0.298) (Figure S3). Overall, narwhals exhibited a greater *F*_ROH_ (mean = 0.033, Q1 = 0.028, Q3 = 0.037) compared to the *F*_ROH_ in bowhead whales (mean = 0.019, Q1 = 0.017, Q3 = 0.022), an indicator of a higher genetic diversity in bowhead whales (Figure 3c, Figure 3d). When examining the number and total lengths of ROHs, we found that narwhals exhibited a greater proportion of shorter ROHs (Figure 3a), which could be a signal of older ROHs and a historic bottleneck [46]. Among narwhal, the northern Hudson Bay subgroup exhibited slightly higher ROH compared to western and eastern Baffin Island narwhals (Figure 3c). This suggests narwhals from northern Hudson Bay may have elevated levels of inbreeding compared to narwhals from Baffin Island.

**Figure 3.**
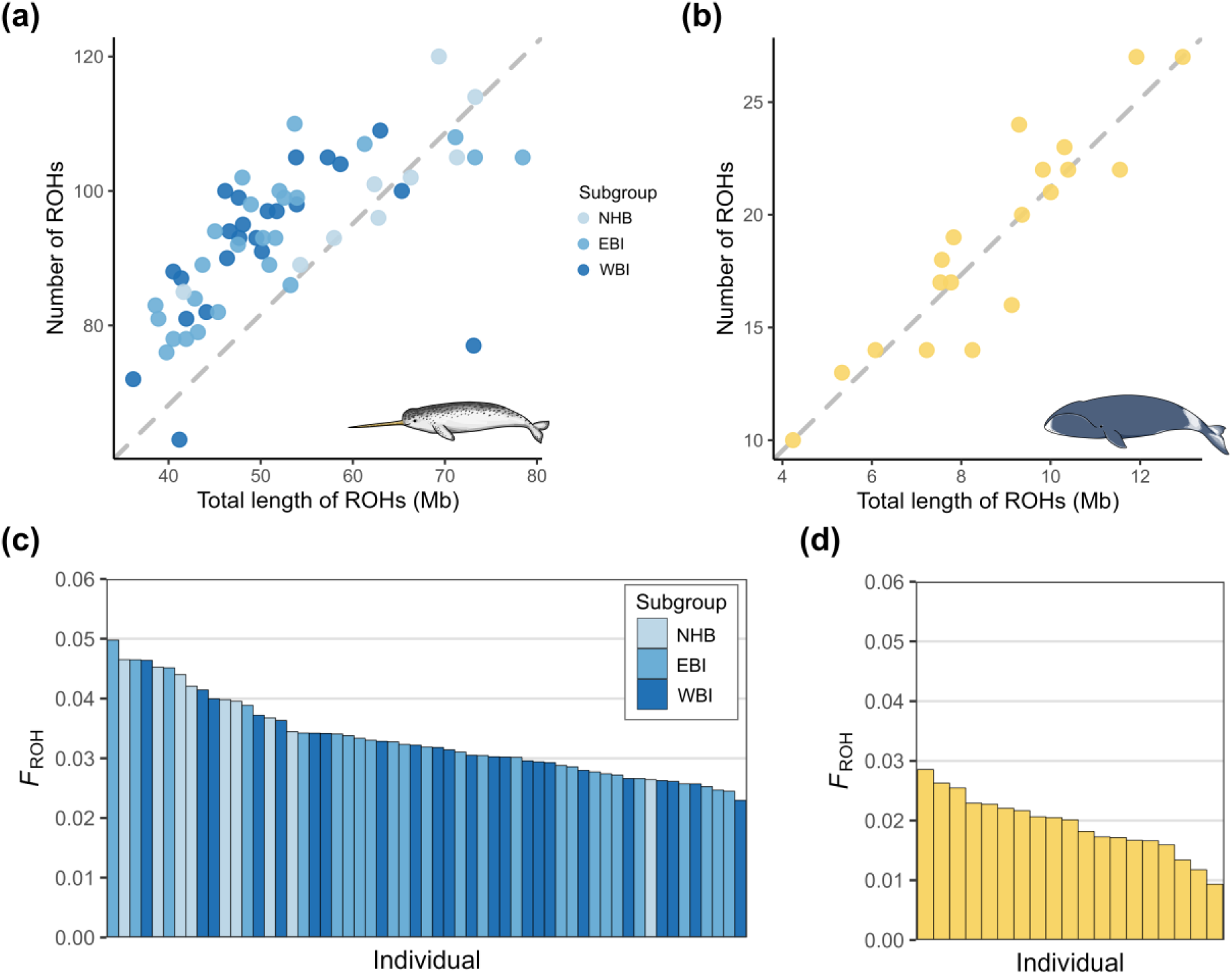
(a) Shorter runs of homozygosity (ROH) in narwhal compared to (b) in bowhead whale. We used the largest scaffolds with even SNP densities to minimize potential biases from genome fragmentation in order to compare species (largest 12 scaffolds in narwhal, largest 200 scaffolds in bowhead). Each point represents an individual and the dashed grey reference line indicates hypothetical symmetry between length and number of ROHs in each species. (c) Proportion of runs of homozygosity (*F*_ROH_) for each narwhal individual, where subgroups are labeled as northern Hudson Bay (NHB), eastern Baffin Island (EBI), and western Baffin Island (WBI), and (d) *F*_ROH_ for each bowhead whale individual.

### (d) Effective population size

Narwhals had an increase in *N*_e_ within the last several thousand years, suggesting a recent population expansion (Figure 4a). When examining narwhal subgroups separately, *N*_e_ was either stable or increasing (Figure S4). Bowhead whales, however, exhibited an increase in *N*_e_ about 5 – 7.5 thousand years ago, then sharply dropped in the last four generations at approximately 200 years ago (Figure 4b), coinciding with commercial whaling.

**Figure 4.**
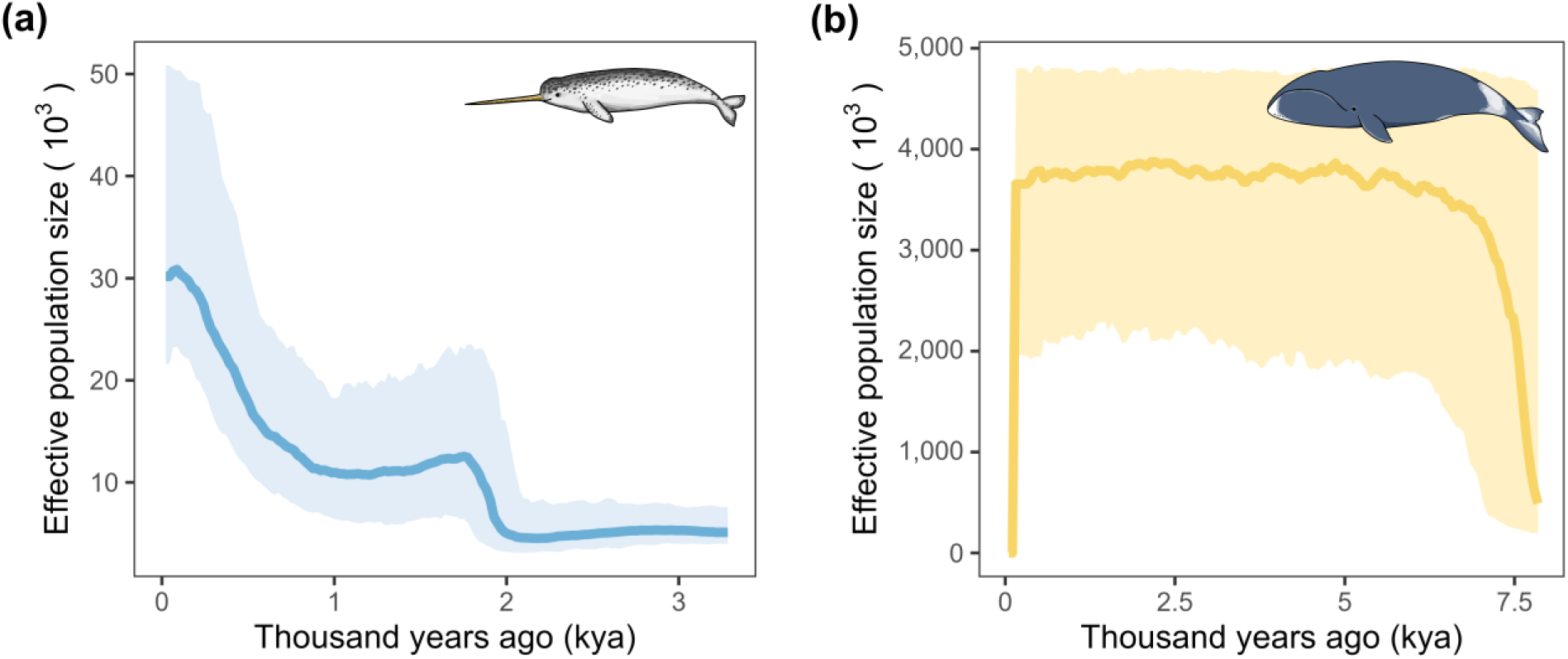
Demographic history within the last 150 generations from GONE analyses show (a) an increase in narwhal effective population size and (b) a sharp decline in bowhead whale effective population size. The median result from 500 iterations is shown by solid lines and 95% confidence intervals shown in lighter shading.

The contemporary *N*_e_ estimate with all narwhal individuals representing the eastern Canadian Arctic as one population was 34,838 (95% CI: 26,720 – 50,037) (Table1). We also estimated contemporary *N*_e_ separately in each narwhal subgroup: western Baffin Island *N*_e_ = 3,620 (95% CI: 3,337 – 3,955); eastern Baffin Island *N*_e_ = 5,587 (95% CI: 4,993 – 6,341); northern Hudson Bay *N*_e_ = 395 (95% CI: 384 – 407) (Table 1). The contemporary *N*_e_ of the Canadian Arctic bowhead whales was 808 (95% CI: 789 – 827) (Table 1). The *N*_e_ /*N* ratio of narwhal was 0.216 as a whole population and 0.021 for the northern Hudson Bay subgroup. The *N*_e_ /*N* ratio of the bowhead whales was 0.125.

**Table 1.**
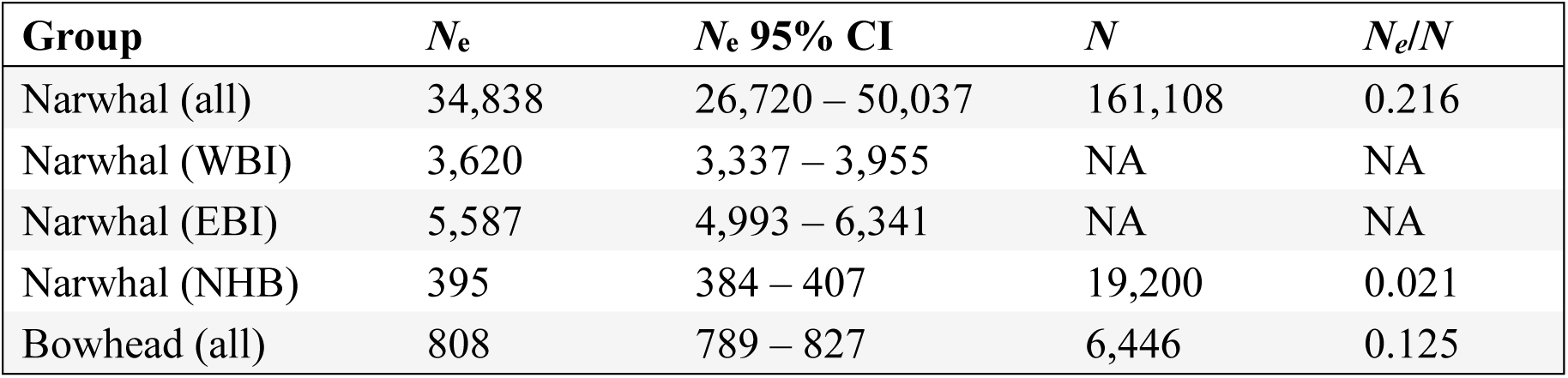
Contemporary effective population sizes (*N*_e_) of eastern Canadian Arctic narwhals and bowhead whales including 95% confidence intervals. Census size from published survey counts (*N*) are included to evaluate *N*_e_/*N* ratios. Groups names that include “all” represent estimates across eastern Canadian Arctic. For narwhal, subgroups are divided as western Baffin Island (WBI), eastern Baffin Island (EBI), and northern Hudson Bay (NHB).

### (e) Historic changes in effective population size

Narwhal and bowhead whale exhibited similar demographic histories throughout the Pleistocene era. Both displayed a steep decline in *N*_e_ approximately 2.5 million years ago (Figure 5), aligning with the onset of the Quaternary Ice Age. In addition, both species exhibited an increase through the last glacial period (11.7 – 115 thousand years ago) followed by a stable *N*_e_ (Figure 5). Iterations of *SMC++* and *PSMC* models for both species are shown in Figures S5, S6, S7, and S8.

**Figure 5.**
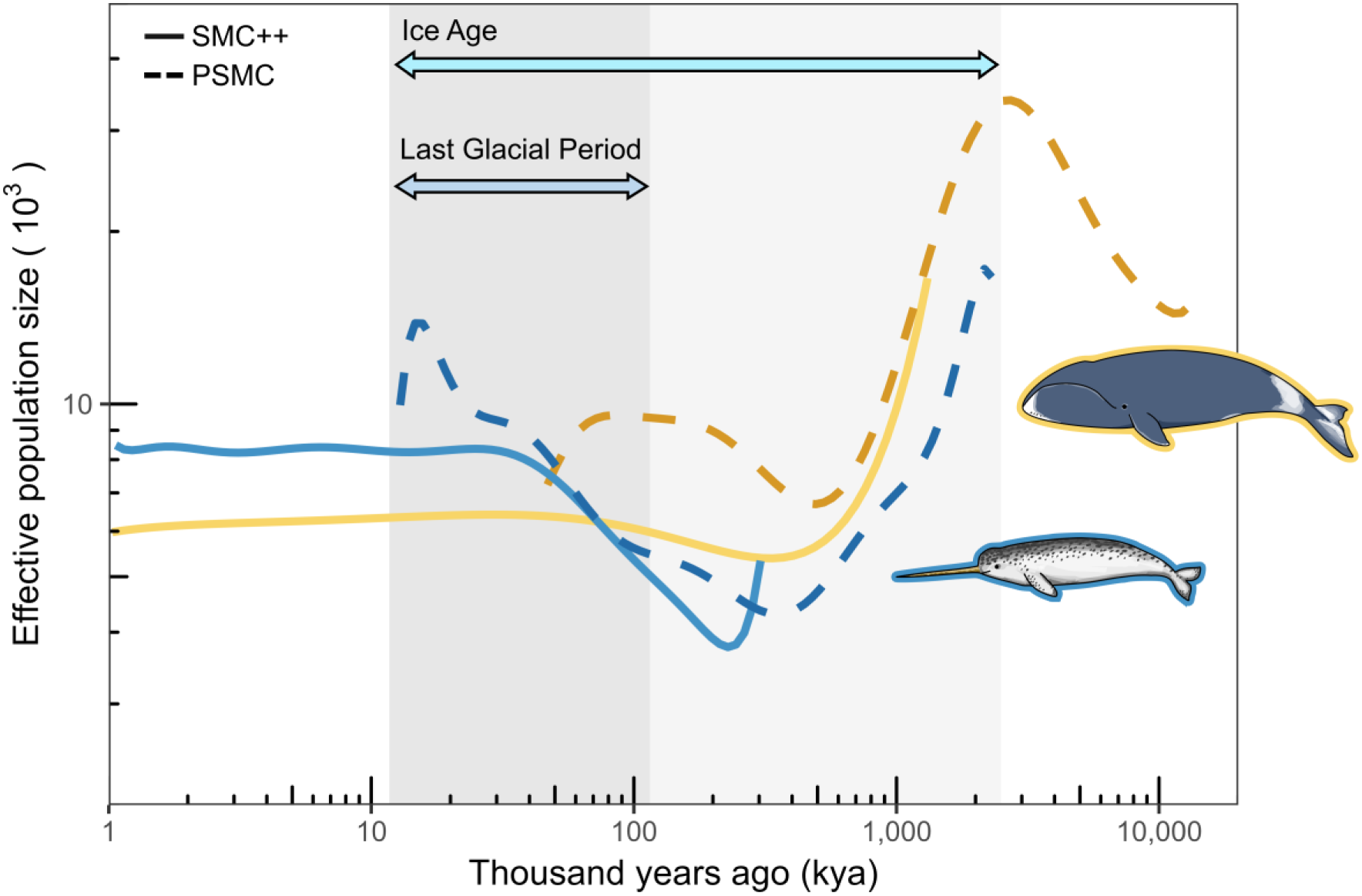
Demographic history of deep past with PSMC (dashed lines) and SMC++ (solid lines). Blue lines represent narwhal results and yellow lines represent results from the bowhead whale. The axes are log-scaled for visualizing trends across a long period. The last Ice Age (11.7 kya -2.5 million years ago) is marked by light and dark grey shading, and the last glacial period (11.7 - 115 kya) is marked by dark gray shading.

## 4. Discussion

### (a) Demographic history and whaling

The recent demographic histories of eastern Canadian Arctic bowhead whales and narwhals reflect the intensity of their history of whaling. Intense commercial whaling significantly decreased bowhead whale *N*_e_ within the past several generations, whereas the minimal opportunistic commercial harvest on narwhal *N*_e_ appears to have had few genetic consequences. Our observation that bowhead whales currently have similar genetic diversity to narwhals demonstrates how the intersection of evolutionary history and whaling influence conservation-oriented thinking in the present day. While narwhals exhibited historically lower genetic diversity, bowhead whales, by contrast, would have had much more genetic diversity and higher *N*_e_ prior to commercial whaling despite now having similar genetic diversity. Bowhead whales’ long generation times and requirement for a larger habitat indicates that commercial whaling may yet have long-lasting effects on genetic variation in the species in the future. This underscores the importance of considering demographic history in population assessments together with genetic diversity [60,61]. A historic population bottleneck in narwhals may have led to their low genetic diversity, indicated by the high frequency of short ROHs, followed by populations persisting at low levels of genetic diversity across a long period of time [59]. Given the increasing *N_e_* of narwhals and their large population size, their low genetic diversity does not suggest concern [60] despite being slightly lower than observed in bowhead whales.

The dramatic decline between previous and current *N*_e_ in bowhead whales could have implications for their genetic health, adaptive potential, and management actions. Although bowhead whale census numbers have increased after commercial whaling ceased, there is evidence it may be plateauing below the pre-commercial whaling carrying capacity estimate [62]. Additionally, populations experience drift at their *N*_e_ and the recovery of genetic diversity will lag behind the recovery of alleles. Because the full effects of drift have not yet been realized, aggressive conservation targets aimed at attaining large population sizes can serve to limit genetic diversity losses. Continued monitoring is necessary to assess for vulnerability to threats such as inbreeding and to provide context for subsistence harvest limits. This present work with Canadian Arctic individuals is an example of how other bowhead whale populations’ trajectories are likely to look given similar levels of commercial whaling. Recent studies in other baleen whales have also documented declines in *N*_e_ coinciding with industrial whaling, such as Atlantic right whales (*Eubalaena glacialis, Eubalaena australis*) [2] and North Atlantic fin whales (*Balaenoptera physalus physalus*) [3].

### (b) Historic demographic trends

Estimating changes in genetic diversity in the two Arctic whales’ history revealed shared trajectories of *N*_e_ through the Pleistocene. Declines in *N*_e_ in narwhal and bowhead whales approximately 2.5 million years ago coincides with an hypothesized supernova explosion that may have contributed to a marine megafauna extinction event and the initiation of the Quaternary Ice Age [63,64]. The cosmic rays from the explosion are believed to have damaged the Earth’s ozone layer [63] and dramatically reduced the Earth’s temperature, subsequently causing major glaciations [65]. While the correlation of the megafauna extinction event and historic declining trend in *N*_e_ in narwhal and bowhead whale is speculative, it is interesting to observe such an impact on the demographic history around a large-scale event that led to other species’ extinctions. Demographic histories of several other whale species also show a similarly timed decline (e.g., fin whale, *Balaenoptera physalus*, [66]; killer whale, *Orcinus orca*, [67]). The long-term decline in *N*_e_ over hundreds of thousands of years in Arctic whales was also observed in Westbury et al. [59] and Cerca et al. [68], and was followed by an increase in *N*_e_ through the last glacial period. The population expansion during the last glacial period could be related to the Arctic whales’ affinity for sea ice, where an expansion of ice sheets may have provided an increase in favourable habitats after intense climatic fluctuations earlier in the Quaternary Ice Age.

### (c) Population structure

Understanding the genetic background of Canadian Arctic narwhal and bowhead whale populations can help guide effective conservation management of these two species. Although the narwhal exhibits low overall genetic variation and diversity [59,69], population substructure in this species may have implications for defining genetic units in the Canadian Arctic. Currently, narwhal are assessed as two genetic populations in Canada, Baffin Bay (Baffin Island) and northern Hudson Bay [70]. Here, we suggest there are three genetically differentiated subgroups in the eastern Canadian Arctic: western Baffin Island, eastern Baffin Island, and northern Hudson Bay. While Baffin Island has been considered a population on its own, previous genetic studies suggested differentiation between Aujuittuq (Grise Fiord) from other parts of Baffin Island [71,72]. Our findings expand this distinction with evidence of substructure between western and eastern Baffin Island including more sublocations, demonstrating that the genetic structure within Baffin Island may be more nuanced than previously thought. Genetic independence may be conserved in the two groups we identified in Baffin Island because of geographically independent wintering grounds which have been identified in satellite telemetry studies [73]. Because of limited sample sizes and minimal genetic differences found in previous work, the conservation status of western and eastern Baffin Island subgroups of narwhals have not been assessed independently [70]. However, Canadian Arctic narwhals have been designated separate management stocks [74].Conservation assessments that consider their genetic segregation in the future would help with effective management strategies. If not assessed individually, at a minimum, our results suggest that the division of Baffin Island narwhal into western and eastern subgroups warrants further study. Interestingly, samples from Kugaaruk and Igloolik grouped with other narwhals from eastern Baffin Island. The level of admixture suggests these sites could be an area of higher gene flow between western and eastern Baffin Island compared to other sites, however, greater sample sizes from Kugaaruk and Igloolik would help investigate this further. Mixing that does occur within Baffin Bay could be contributed by narwhals mating during migration. Mating is thought to occur mostly between narwhals migrating to the same summering area [75]; however, the timing of mating in narwhals varies, spanning February to April [76] and possibly between May and June [77]. Despite weak values of genetic differentiation (limited structuring also observed in mitogenomes [69]), overall these results demonstrate a lack of panmixia, and present subtle genetic differentiation positively correlated with increasing geographical distance.

The northern Hudson Bay subpopulation warrants conservation priority within Canadian Arctic narwhals in the context of genetic diversity, due to greater levels of inbreeding, and lower effective population size. Aerial survey abundance estimates suggest that the northern Hudson Bay narwhal population is increasing, albeit with slowing growth rate as it reaches carrying capacity [78]. At this slowed growth rate Biddlecombe and Watt [78] predicted that even modest harvests could result in a decline in abundance. Our low estimates of *N*_e_ also suggest this population may be vulnerable to harvest and highlight the need to consider both abundance and genetic population structure and genetic resilience when assessing management and conservation strategies [79].

In bowhead whales, the lack of clear structuring is consistent with high gene flow and limited structure across the eastern Canadian Arctic [80,81]. We would expect that the broad geographical range of sample collection would capture evidence of clustering if there were distinct genetic populations within the Canadian Arctic, however, sample sizes may limit our inferences of population structure. While outliers in the PCA could represent migrants from other genetic populations, more samples would help investigate their structure in more detail. Within large geographical regions, it appears gene flow is generally high in bowhead whale groups (within the Canadian Arctic [28], East Greenland [30], and western Arctic [82]). Bowhead whales are capable of large movements [83] and separate populations have overlapped summer distributions.

### (d) Conclusions

The eastern Canadian Arctic bowhead whale’s sharp decline in *N*_e_ within the past several generations demonstrate how overharvest associated with commercial whaling can significantly alter genetic dynamics of populations. Despite similar metrics of genetic diversity in bowhead whale compared to narwhal, bowhead whale’s recent sharp decline should lead to continued effects of genetic drift in the future. Additionally, both species exhibited overall low genomic variation, with subtle population differentiation in narwhal suggesting the presence of three Canadian Arctic narwhal genetic subgroups. Together, these results provide a deeper understanding of endemic whale populations in the Canadian Arctic, providing critical background information for guiding conservation management strategies.

## Acknowledgements

We would like to thank the Greenland Institute of Natural Resources, the Newfoundland and Labrador Whale Release and Strandings Network, local communities, and the following Hunters and Trappers Organizations/Associations/Committees for collecting samples in this study: Aiviit, Aiviq, Amarok, Arviq, Nangmautaq, Gjoa Haven, Igloolik, Ikajutit, Iviq, Kaniqliniq, Kaniqsujuaq, Kurtairujuark, Mittimatalik, Nattivak, Pangnirtung, Resolute Bay, Sachs Harbour, Sanirajak, Spence Bay, Tuktoyaktuk. We are grateful to the Hospital for Sick Children (SickKids) for sequencing our DNA samples. This research was enabled in part by support provided by the Prairies DRI and the Digital Research Alliance of Canada (alliancecan.ca).

## Data accessibility statement

Raw data are uploaded on NCBI BioProject PRJNA1026538 (narwhal) and BioProject PRJNA1026863 (bowhead whale). Code for all analyses are available at github.com/edegreef/arctic-whales-resequencing.

## References

1. Estes JA, Demaster DP, Doak DF, Williams TM, Brownell RL. 2006 Whales, Whaling, and Ocean Ecosystems. 1st edn. University of California Press. See https://www.jstor.org/stable/10.1525/j.ctt1ppsvh.

2. Crossman CA, Fontaine MC, Frasier TR. 2023 A comparison of genomic diversity and demographic history of the North Atlantic and Southwest Atlantic southern right whales. Molecular Ecology **n/a**. (doi:10.1111/mec.17099)

3. Wolf M, de Jong M, Halldórsson SD, Árnason Ú, Janke A. 2022 Genomic Impact of Whaling in North Atlantic Fin Whales. Molecular Biology and Evolution 39, msac094. (doi:10.1093/molbev/msac094)

4. Ross W. 1993 Commercial whaling in the North Atlantic sector. In J.J. Burns, J.J. Montague, C.J. Cowles (Eds.), The Bowhead Whale, Society for Marine Mammalogy Special Publication Number 2, pp. 511–561. Lawrence, Kansas.

5. Rocha, Jr. RC, Clapham PJ, Ivashchenko Y. 2015 Emptying the Oceans: A Summary of Industrial Whaling Catches in the 20th Century. MFR 76, 37–48. (doi:10.7755/MFR.76.4.3)

6. Lynch M. 2010 Evolution of the mutation rate. Trends Genet 26, 345–352. (doi:10.1016/j.tig.2010.05.003)

7. Kardos M et al. 2023 Inbreeding depression explains killer whale population dynamics. Nat Ecol Evol (doi:10.1038/s41559-023-01995-0)

8. Agrelo M et al. 2021 Ocean warming threatens southern right whale population recovery. Science Advances 7, eabh2823. (doi:10.1126/sciadv.abh2823)

9. Meyer-Gutbrod EL, Greene CH. 2018 Uncertain recovery of the North Atlantic right whale in a changing ocean. Global Change Biology 24, 455–464. (doi:10.1111/gcb.13929)

10. Tulloch VJD, Plagányi ÉE, Brown C, Richardson AJ, Matear R. 2019 Future recovery of baleen whales is imperiled by climate change. Global Change Biology 25, 1263–1281. (doi:10.1111/gcb.14573)

11. Rantanen M, Karpechko AY, Lipponen A, Nordling K, Hyvärinen O, Ruosteenoja K, Vihma T, Laaksonen A. 2022 The Arctic has warmed nearly four times faster than the globe since 1979. Commun Earth Environ 3, 1–10. (doi:10.1038/s43247-022-00498-3)

12. Chambault P et al. 2020 The impact of rising sea temperatures on an Arctic top predator, the narwhal. Sci Rep 10, 18678. (doi:10.1038/s41598-020-75658-6)

13. Simmonds MP, Isaac SJ. 2007 The impacts of climate change on marine mammals: early signs of significant problems. Oryx 41, 19–26. (doi:10.1017/S0030605307001524)

14. Flores H et al. 2023 Sea-ice decline could keep zooplankton deeper for longer. Nat. Clim. Chang. 13, 1122–1130. (doi:10.1038/s41558-023-01779-1)

15. Florko KRN, Tai TC, Cheung WWL, Ferguson SH, Sumaila UR, Yurkowski DJ, Auger-Méthé M. 2021 Predicting how climate change threatens the prey base of Arctic marine predators. Ecology Letters 24, 2563–2575. (doi:10.1111/ele.13866)

16. Higdon JW, Ferguson SH. 2009 Loss of Arctic sea ice causing punctuated change in sightings of killer whales (*Orcinus orca*) over the past century. Ecological Applications 19, 1365–1375. (doi:10.1890/07-1941.1)

17. Mudryk LR, Dawson J, Howell SEL, Derksen C, Zagon TA, Brady M. 2021 Impact of 1, 2 and 4 °C of global warming on ship navigation in the Canadian Arctic. Nat. Clim. Chang. 11, 673–679. (doi:10.1038/s41558-021-01087-6)

18. Lowry L, Laidre K, Reeves R. 2017 Monodon monoceros. The IUCN Red List of Threatened Species 2017. IUCN Red List of Threatened Species. See https://www.iucnredlist.org/species/13704/50367651 (accessed on 23 October 2023).

19. Ross W, McIver A. 1982 Distribution of the kills of bowhead whales and other sea mammals by Davis Strait whalers, 1820, to 1910.

20. Stewart DB. 2008 Commercial and subsistence harvests of narwhal (Monodon monoceros) from the Canadian eastern Arctic., ii + 97 p.

21. McLeod BA, Brown MW, Moore MJ, Stevens W, Barkham SH, Barkham M, White BN. 2009 Bowhead whales, and not right whales, were the primary target of 16th- to 17th-century Basque Whalers in the Western North Atlantic. ARCTIC 61, 61. (doi:10.14430/arctic7)

22. Higdon JeffW. 2010 Commercial and subsistence harvests of bowhead whales (Balaena mysticetus) in eastern Canada and West Greenland. JCRM 11, 185–216. (doi:10.47536/jcrm.v11i2.623)

23. COSEWIC. 2009 COSEWIC assessment and update status report on the Bowhead Whale Balaena mysticetus, Bering-Chukchi-Beaufort population and Eastern Canada-West Greenland population, in Canada., vii + 49 pp.

24. Garde E, Hansen SH, Ditlevsen S, Tvermosegaard KB, Hansen J, Harding KC, Heide-Jørgensen MP. 2015 Life history parameters of narwhals (*Monodon monoceros*) from Greenland. JMAMMA 96, 866–879. (doi:10.1093/jmammal/gyv110)

25. Keane M et al. 2015 Insights into the Evolution of Longevity from the Bowhead Whale Genome. Cell Reports 10, 112–122. (doi:10.1016/j.celrep.2014.12.008)

26. Hohenlohe PA, Funk WC, Rajora OP. 2021 Population genomics for wildlife conservation and management. Molecular Ecology 30, 62–82. (doi:10.1111/mec.15720)

27. Waples RS. 2022 What Is *N* e, Anyway? Journal of Heredity 113, 371–379. (doi:10.1093/jhered/esac023)

28. Wang J, Santiago E, Caballero A. 2016 Prediction and estimation of effective population size. Heredity 117, 193–206. (doi:10.1038/hdy.2016.43)

29. Bolger AM, Lohse M, Usadel B. 2014 Trimmomatic: a flexible trimmer for Illumina sequence data. Bioinformatics 30, 2114–2120. (doi:10.1093/bioinformatics/btu170)

30. Damas J et al. 2022 Evolution of the ancestral mammalian karyotype and syntenic regions. Proceedings of the National Academy of Sciences 119, e2209139119. (doi:10.1073/pnas.2209139119)

31. Li H, Durbin R. 2009 Fast and accurate short read alignment with Burrows–Wheeler transform. Bioinformatics 25, 1754–1760. (doi:10.1093/bioinformatics/btp324)

32. Broad Institute. 2019 Picard Toolkit. GitHub Repository. See https://broadinstitute.github.io/picard/ (accessed on 15 August 2023).

33. Li H et al. 2009 The Sequence Alignment/Map format and SAMtools. Bioinformatics 25, 2078–2079. (doi:10.1093/bioinformatics/btp352)

34. McKenna A et al. 2010 The Genome Analysis Toolkit: a MapReduce framework for analyzing next-generation DNA sequencing data. Genome Res 20, 1297–1303. (doi:10.1101/gr.107524.110)

35. Rimmer A, Phan H, Mathieson I, Iqbal Z, Twigg SRF, WGS500 Consortium, Wilkie AOM, McVean G, Lunter G. 2014 Integrating mapping-, assembly- and haplotype-based approaches for calling variants in clinical sequencing applications. Nat Genet 46, 912–918. (doi:10.1038/ng.3036)

36. Danecek P et al. 2011 The variant call format and VCFtools. Bioinformatics 27, 2156–2158. (doi:10.1093/bioinformatics/btr330)

37. Purcell S et al. 2007 PLINK: a tool set for whole-genome association and population-based linkage analyses. Am J Hum Genet 81, 559–575. (doi:10.1086/519795)

38. Privé F, Luu K, Vilhjálmsson BJ, Blum MGB. 2020 Performing Highly Efficient Genome Scans for Local Adaptation with R Package pcadapt Version 4. Molecular Biology and Evolution 37, 2153–2154. (doi:10.1093/molbev/msaa053)

39. Frichot E, François O. 2015 LEA: An R package for landscape and ecological association studies. Methods in Ecology and Evolution 6, 925–929. (doi:10.1111/2041-210X.12382)

40. R Core Team. 2022 R: A language and environment for statistical computing.

41. Reich D, Thangaraj K, Patterson N, Price AL, Singh L. 2009 Reconstructing Indian population history. Nature 461, 489–494. (doi:10.1038/nature08365)

42. Pante E, Simon-Bouhet B. 2013 marmap: A Package for Importing, Plotting and Analyzing Bathymetric and Topographic Data in R. PLOS ONE 8, e73051. (doi:10.1371/journal.pone.0073051)

43. Dray S, Dufour A-B. 2007 The ade4 Package: Implementing the Duality Diagram for Ecologists. Journal of Statistical Software 22, 1–20. (doi:10.18637/jss.v022.i04)

44. Reed DH, Frankham R. 2003 Correlation between Fitness and Genetic Diversity. Conservation Biology 17, 230–237. (doi:10.1046/j.1523-1739.2003.01236.x)

45. Goudet J. 2005 hierfstat, a package for r to compute and test hierarchical F-statistics. Molecular Ecology Notes 5, 184–186. (doi:10.1111/j.1471-8286.2004.00828.x)

46. Ceballos FC, Joshi PK, Clark DW, Ramsay M, Wilson JF. 2018 Runs of homozygosity: windows into population history and trait architecture. Nat Rev Genet 19, 220–234. (doi:10.1038/nrg.2017.109)

47. Foote AD et al. 2021 Runs of homozygosity in killer whale genomes provide a global record of demographic histories. Molecular Ecology 30, 6162–6177. (doi:10.1111/mec.16137)

48. Santiago E, Novo I, Pardiñas AF, Saura M, Wang J, Caballero A. 2020 Recent Demographic History Inferred by High-Resolution Analysis of Linkage Disequilibrium. Molecular Biology and Evolution 37, 3642–3653. (doi:10.1093/molbev/msaa169)

49. Taylor B, Chivers S, Larese J, Perrin W. 2007 Generation length and percent mature estimates for IUCN assessments of cetaceans. NOAA, NMFS, Southwest Fisheries Science Center Administrative Report LJ-07-01

50. Archer FI, Adams PE, Schneiders BB. 2017 stratag: An r package for manipulating, summarizing and analysing population genetic data. Molecular Ecology Resources 17, 5–11. (doi:10.1111/1755-0998.12559)

51. Waples RS, Do C. 2008 ldne: a program for estimating effective population size from data on linkage disequilibrium. Molecular Ecology Resources 8, 753–756. (doi:10.1111/j.1755-0998.2007.02061.x)

52. Waples RS, Grewe PM, Bravington MW, Hillary R, Feutry P. 2018 Robust estimates of a high Ne/N ratio in a top marine predator, southern bluefin tuna. Science Advances 4, eaar7759. (doi:10.1126/sciadv.aar7759)

53. Doniol-Valcroze T, Gosselin J-F, Pike DG, Lawson JW, Asselin NC, Hedges K, Ferguson SH. 2020 Narwhal Abundance in the Eastern Canadian High Arctic in 2013. NAMMCOSP 11. (doi:10.7557/3.5100)

54. DFO. 2020 Abundance Estimate of the Northern Hudson Bay Narwhal Population from the 2018 Aerial Survey. Canadian Science Advisory Secretariat

55. Doniol-Valcroze T, Gosselin J-F, Pike DG, Lawson JW, Asselin NC, Hedges KJ, Ferguson SH. 2019 Distribution and Abundance of the Eastern Canada – West Greenland Bowhead Whale Population Based on the 2013 High Arctic Cetacean Survey. NAMMCO Scientific Publications 11. (doi:10.7557/3.5315)

56. Nadachowska-Brzyska K, Konczal M, Babik W. 2022 Navigating the temporal continuum of effective population size. Methods in Ecology and Evolution 13, 22–41. (doi:10.1111/2041-210X.13740)

57. Li H, Durbin R. 2011 Inference of human population history from individual whole-genome sequences. Nature 475, 493–496. (doi:10.1038/nature10231)

58. Terhorst J, Kamm JA, Song YS. 2017 Robust and scalable inference of population history from hundreds of unphased whole genomes. Nat Genet 49, 303–309. (doi:10.1038/ng.3748)

59. Westbury MV, Petersen B, Garde E, Heide-Jørgensen MP, Lorenzen ED. 2019 Narwhal Genome Reveals Long-Term Low Genetic Diversity despite Current Large Abundance Size. iScience 15, 592–599. (doi:10.1016/j.isci.2019.03.023)

60. Schmidt C, Hoban S, Hunter M, Paz-Vinas I, Garroway CJ. 2023 Genetic diversity and IUCN Red List status. Conservation Biology 37, e14064. (doi:10.1111/cobi.14064)

61. Wootton E, Robert C, Taillon J, Côté SD, Shafer ABA. 2023 Genomic health is dependent on long-term population demographic history. Molecular Ecology 32, 1943–1954. (doi:10.1111/mec.16863)

62. Biddlecombe BA, Ferguson SH, Heide-Jørgensen MP, Gillis DM, Watt CA. 2023 Estimating abundance of Eastern Canada-West Greenland bowhead whales using genetic mark-recapture analyses. Global Ecology and Conservation 45, e02524. (doi:10.1016/j.gecco.2023.e02524)

63. Benítez N, Maíz-Apellániz J, Canelles M. 2002 Evidence for Nearby Supernova Explosions. Phys. Rev. Lett. 88, 081101. (doi:10.1103/PhysRevLett.88.081101)

64. Melott AL, Marinho F, Paulucci L. 2019 Hypothesis: Muon Radiation Dose and Marine Megafaunal Extinction at the End-Pliocene Supernova. Astrobiology 19, 825–830. (doi:10.1089/ast.2018.1902)

65. Korschinek G. 2017 Mass extinctions and supernova explosions. pp. 2419–2430. (doi:10.1007/978-3-319-21846-5_22)

66. Árnason Ú, Lammers F, Kumar V, Nilsson MA, Janke A. 2018 Whole-genome sequencing of the blue whale and other rorquals finds signatures for introgressive gene flow. Science Advances 4, eaap9873. (doi:10.1126/sciadv.aap9873)

67. Moura AE et al. 2014 Population genomics of the killer whale indicates ecotype evolution in sympatry involving both selection and drift. Mol Ecol 23, 5179–5192. (doi:10.1111/mec.12929)

68. Cerca J, Westbury MV, Heide-Jørgensen MP, Kovacs KM, Lorenzen ED, Lydersen C, Shpak OV, Wiig Ø, Bachmann L. 2022 High genomic diversity in the endangered East Greenland Svalbard Barents Sea stock of bowhead whales (Balaena mysticetus). Sci Rep 12, 6118. (doi:10.1038/s41598-022-09868-5)

69. Louis M et al. 2020 Influence of past climate change on phylogeography and demographic history of narwhals, Monodon monoceros. Proc. R. Soc. B. 287, 20192964. (doi:10.1098/rspb.2019.2964)

70. COSEWIC. 2004 COSEWIC assessment and update status report on the narwhal Monodon monoceros in Canada.

71. de March BGE, Tenkula DA, Postma LD. 2003 Molecular genetics of narwhal (Monodon monoceros) from Canada and West Greenland (1982-2001).

72. Petersen SD, Tenkula D, Ferguson SH. 2011 Population genetic structure of narwhal (Monodon monoceros).

73. Laidre K, Heide-Jørgensen M, Dietz R, Hobbs R, Jørgensen O. 2003 Deep-diving by narwhals Monodon monoceros: differences in foraging behavior between wintering areas? Mar. Ecol. Prog. Ser. 261, 269–281. (doi:10.3354/meps261269)

74. DFO. 2015 Abundance estimates of narwhal stocks in the Canadian High Arctic in 2013. Canadian Science Advisory Secretariat Science Advisory Report 2015/046.

75. Heide-Jørgensen MP, Richard PR, Dietz R, Laidre KL. 2013 A metapopulation model for Canadian and West Greenland narwhals. Animal Conservation 16, 331–343. (doi:10.1111/acv.12000)

76. Best RC, Fisher HD. 1974 Seasonal breeding of the narwhal (Monodon monoceros L.). Can. J. Zool. 52, 429–431. (doi:10.1139/z74-052)

77. Heide-Jørgensen MP, Garde E. 2011 Fetal growth of narwhals (Monodon monoceros). Marine Mammal Science 27, 659–664. (doi:10.1111/j.1748-7692.2010.00423.x)

78. Biddlecombe BA, Watt CA. 2022 Modeling population trajectory and probability of decline in northern Hudson Bay narwhals (*Monodon monoceros*). Marine Mammal Science 38, 1357–1370. (doi:10.1111/mms.12925)

79. Taylor BL. 2005 Identifying units to conserve using genetic data. In J.E.I. Reynolds, W.F. Perrin, R. R. Reeves, S. Montgomery, T.J. Ragen (Eds.), Mar. Mammal. Res. Conserv. Crisis, pp. 165–176. Baltimore, MD: Johns Hopkins University Press.

80. Elizabeth Alter S et al. 2012 Gene flow on ice: the role of sea ice and whaling in shaping Holarctic genetic diversity and population differentiation in bowhead whales (*Balaena mysticetus*). Ecol Evol 2, 2895–2911. (doi:10.1002/ece3.397)

81. Postma LD, Dueck LP, Heide-Jørgensen MP. 2005 Comparaison de la génétique moléculaire des baleines boréales (Balaena mysticetus) des eaux de l’est de l’Arctique canadien et de l’ouest du Groenland. Canadian Science Advisory Secretariat Research Document 2005/004, 29.

82. Givens GH, Huebinger RM, Patton JC, Postma LD, Lindsay M, Suydam RS, George JC, Matson CW, Bickham JW. 2010 Population Genetics of Bowhead Whales (Balaena mysticetus) in the Western Arctic. Arctic 63, 1–12.

83. Chambault P, Albertsen CM, Patterson TA, Hansen RG, Tervo O, Laidre KL, Heide-Jørgensen MP. 2018 Sea surface temperature predicts the movements of an Arctic cetacean: the bowhead whale. Sci Rep 8, 9658. (doi:10.1038/s41598-018-27966-1)

