## Supplementary Material for "Unraveling the genetic legacy of commercial whaling in bowhead whales and narwhals"

### Methods

*Identifying sex-linked scaffolds*

We determined sex-linked scaffolds through a coverage comparison method using *DifCover* [1] and the *SexFindR* pipeline [2]. Given that females have two X chromosomes and males have an X and Y chromosome, we identified which scaffolds had approximately double or half of the coverages when comparing male and female samples. For both the narwhal and bowhead whale, we selected four samples with previously known sexes (two males and two females) and completed coverage comparisons in pairs with each combination (male1 & female1, male1 & female2, male2 & female1, male2 & female2, male1 & male2, female1 & female2). The four male & female combinations were used to identify scaffolds that were notably different, or enriched, in one sex or the other. Same-sex comparisons were used as controls. We analyzed outputs in R version 4.2.1 [3]. We then annotated scaffold windows with *BEDTools* v2.25.0 [4], and then extracted scaffolds that were enriched in the four male & female combinations and not enriched in the controls to identify sex-linked scaffolds. In the narwhal, we used samples ARGF_07_1125 (male1), ARAB_08_1164 (female1), ARIQ_DFO_11_1071 (male2), and ARBI_05_1092 (female2). For the bowhead whale, we used samples BM_NSA_2008_02 (male1), BM_NSA_2009_02 (female1), BM_NSA_2010_01 (male2), and BM_NSA_2012_02 (female2).

*Demographic history - PSMC*

We used *PSMC*, a pairwise sequentially Markovian coalescent [5] to explore changes in genetic diversity in the deep past. Here, we used one high-coverage sample for each species using sequence data prior to down-sampling: ARCR_07_1065 (20x coverage) for the narwhal, and 88_Pang (18x coverage) for the bowhead whale. We called SNPs through the mpileup function (-C50, -Q20, -q20) in *Samtools* v1.9 [6] using the species' respective reference genomes. In the PSMC analyses, we used parameters N=25, t=15, r=5, and p=4+25*2+4+6. In the final plot, we multiplied the number of generations with respective generation times for each species (21.9 years for the narwhal; 52.3 years for the bowhead whale [7]) and used previously documented mutation rates (narwhal $\mu = 1.56 \times 10^{-8}$; bowhead whale $\mu = 2.69 \times 10^{-8}$ [8]).

*Demographic history – SMC++*

For the second demographic history model, we used *SMC++*, a sequentially Markovian coalescent analysis that combines site frequency spectrum and linkage information [9]. *SMC++* takes in multiple genomes and improves accuracy compared to previous SMC methods for inferring $N_e$ in recent past. We used autosomal SNPs and excluded sites out of Hardy-Weinberg equilibrium (heterozygous frequency threshold > 0.6). To distinguish missing regions from long runs of homozygosity, we used indels and unused loci to create a masked file with *bedops* [10], which we included into the SMC++ model. When creating composite likelihoods (through smc++ vcf2smc) for each population, we set individual(s) for a distinguished lineage. For the narwhal data as one population we used individuals 94_RAHM_IQ_152 and ARGF_01_1068 as distinguished lineages. To explore potential differences between narwhal subgroups, we ran an additional set of models for each subpopulation cluster using individuals ARRB_16_1379 for the northern Hudson Bay subgroup, 94_RAHM_IQ_152 for the eastern Baffin Bay subgroup, and 94_RAHM_IQ_241_GF for the western Baffin Bay subgroup as distinguished lineages. In the bowhead whale, we estimated demographic history as one population and used individuals 88_Pang and WBF_2005_0298 as distinguished lineages. In the SMC++ estimate, we ran 100 iterations for each population using the same mutation rates used in the PSMC model. Model parameters included a regularization penalty of 4.0, a non-segregating site cut-off of 100,000, thinning of 2,000 and timepoints of 10 to 10,000 generations. We used the same generation times as used in the PSMC model.

**References**

1. Smith JJ *et al.* 2018 The sea lamprey germline genome provides insights into programmed genome rearrangement and vertebrate evolution. *Nat Genet* **50**, 270–277. (doi:10.1038/s41588-017-0036-1)

2. Grayson P, Wright A, Garroway CJ, Docker MF. 2022 SexFindR: A computational workflow to identify young and old sex chromosomes. , 2022.02.21.481346. (doi:10.1101/2022.02.21.481346)

3. R Core Team. 2022 R: A language and environment for statistical computing.

4. Quinlan AR, Hall IM. 2010 BEDTools: a flexible suite of utilities for comparing genomic features. *Bioinformatics* **26**, 841–842. (doi:10.1093/bioinformatics/btq033)

5. Li H, Durbin R. 2011 Inference of human population history from individual whole-genome sequences. *Nature* **475**, 493–496. (doi:10.1038/nature10231)

6. Li H *et al.* 2009 The Sequence Alignment/Map format and SAMtools. *Bioinformatics* **25**, 2078–2079. (doi:10.1093/bioinformatics/btp352)

7. Taylor B, Chivers S, Larese J, Perrin W. 2007 Generation length and percent mature estimates for IUCN assessments of cetaceans. *NOAA, NMFS, Southwest Fisheries Science Center Administrative Report LJ-07-01*

8. Westbury MV, Petersen B, Garde E, Heide-Jørgensen MP, Lorenzen ED. 2019 Narwhal Genome Reveals Long-Term Low Genetic Diversity despite Current Large Abundance Size. *iScience* **15**, 592–599. (doi:10.1016/j.isci.2019.03.023)

9. Terhorst J, Kamm JA, Song YS. 2017 Robust and scalable inference of population history from hundreds of unphased whole genomes. *Nat Genet* **49**, 303–309. (doi:10.1038/ng.3748)

10. Neph S *et al.* 2012 BEDOPS: high-performance genomic feature operations. *Bioinformatics* **28**, 1919–1920. (doi:10.1093/bioinformatics/bts277)

**Tables & Figures**

**Table S1**. Site names and number of samples (including kin pairs) collected from each site. The year of sample collection is listed in each species' respective columns.

| **Site name** | **Alternate name** | **Narwhal (*n*)** | **Collection year(s)** | **Bowhead whale (*n*)** | **Collection year(s)** |
|---|---|---|---|---|---|
| Uqsuqtuuq | Gjoa Haven | | | 1 | 2002 |
| Qausuittuq | Resolute Bay | 9 | 2002-2012 | | |
| Taloyoak | Spence Bay | 6 | 1996-2009 | 1 | 2012 |
| Kangiq&iniq | Rankin Inlet | | | 1 | 2009 |
| Kugaaruk | Pelly Bay | 3 | 2020 | 1 | 2011 |
| Naujaat | Repulse Bay | 9 | 1993-2020 | 1 | 2005 |
| Ikpiarjuk | Arctic Bay | 9 | 1987-2010 | 1 | 2012 |
| Salliq | Coral Harbour | | | 2 | 1999-2000 |
| Aujuittuq | Grise Fiord | 10 | 1994-2007 | | |
| Igloolik | | 3 | 1982-2006 | | |
| Sanirajak | Hall Beach | | | 2 | 2008-2020 |
| Kinngait | Cape Dorset | | | 1 | 2009 |
| Mittimatalik | Pond Inlet | 4 | 1992-2017 | 1 | 2010 |
| Kangiqsujuaq | Wakeham Bay | | | 1 | 2009 |
| Kangiqtugaapik | Clyde River | 2 | 2007 | 1 | 2014 |
| Iqaluit | | | | 1 | 2011 |
| Panniqtuuq | Pangnirtung | 1 | 2019 | 2 | 1998 |
| Qikiqtarjuaq | Broughton Island | 6 | 1994-2011 | | |
| West Greenland | | | | 2 | 2007 |
| Newfoundland | | | | 1 | 2005 |
| Greenland | | | | 1 | 2006 |

**Table S2**. Sample information for each narwhal individual, including sample ID, sex determined by genome coverage comparisons, modal coverage, final modal coverages for down-sampled samples, the frequency of SNP missingness, and notes on which individuals were removed for downstream analyses.

| Sample ID | Sex | Modal coverage | Down-sampled modal coverage | SNP miss. freq. | Removed individual (X) |
|---|---|---|---|---|---|
| 94_RAHM_IQ_152 | F | 15 | 10 | 0.025 | |
| 94_RAHM_IQ_162 | M | 18 | 9 | 0.033 | X - duplicate with RA_HM_IQ_121 |
| 94_RAHM_IQ_241_GF | M | 23 | 9 | 0.033 | |
| 94_RAHM_IQ_243_GF | F | 22 | 10 | 0.026 | |
| 94_RAHM_IQ_244_GF | M | 16 | 10 | 0.028 | |
| ARAB_04_03 | M | 16 | 10 | 0.025 | |
| ARAB_04_04 | F | 22 | 10 | 0.024 | |
| ARAB_08_1163 | M | 12 | 10 | 0.026 | |
| ARAB_08_1164 | F | 14 | 10 | 0.023 | |
| ARAB_10_1221 | F | 19 | 10 | 0.024 | |
| ARAB_10_1223 | M | 22 | 10 | 0.030 | |
| ARAB_99_1005 | F | 14 | 10 | 0.026 | |
| ARAB_99_1006 | M | 20 | 9 | 0.034 | |
| ARBI_05_1092 | F | 15 | 10 | 0.027 | |
| ARBI_05_1093 | M | 12 | 10 | 0.030 | |
| ARCR_07_1065 | F | 20 | 10 | 0.024 | |
| ARCR_07_1070 | M | 13 | 9 | 0.030 | |
| ARGF_01_1068 | M | 15 | 10 | 0.026 | |
| ARGF_01_1082 | M | 14 | 10 | 0.028 | |
| ARGF_01_1088 | M | 17 | 10 | 0.029 | |
| ARGF_07_1121 | M | 19 | 10 | 0.026 | |
| ARGF_07_1125 | M | 16 | 10 | 0.028 | |
| ARGF_07_1127 | F | 21 | 10 | 0.025 | |
| ARGF_07_1131 | F | 17 | 10 | 0.023 | |
| ARIG_06_1222 | F | 19 | 10 | 0.025 | |
| ARIQ_DFO_11_1071 | M | 16 | 10 | 0.028 | |
| ARPB_xx_1091 | F | 14 | 10 | 0.025 | |
| ARPB_xx_1095 | M | 9 | | 0.037 | |
| ARPB_xx_1100 | F | 16 | 10 | 0.025 | |
| ARPG_19_1614 | M | 14 | 10 | 0.031 | |
| ARPI_04_1184 | M | 7 | | 0.069 | |
| ARPI_08_0082 | F | 14 | 9 | 0.027 | |
| ARPI_17_1336 | F | 16 | 10 | 0.025 | |
| ARRB_16_1379 | M | 12 | 10 | 0.028 | |
| ARRB_99_1001 | F | 19 | 10 | 0.023 | |
| ARRB_99_1026 | M | 21 | 10 | 0.030 | |
| ARRB_xx_1421 | F | 14 | 10 | 0.023 | |
| ARRB_xx_1457 | M | 16 | 10 | 0.028 | |
| ARRE_02_1094 | M | 12 | 10 | 0.024 | |
| ARRE_02_1106 | M | 20 | 10 | 0.029 | |
| ARRE_06_1144 | M | 18 | 10 | 0.030 | |
| ARRE_06_1146 | M | 24 | 10 | 0.029 | |
| ARRE_06_1157 | M | 23 | 10 | 0.028 | |
| ARRE_06_1158 | M | 11 | 10 | 0.030 | |
| ARRE_06_1159 | F | 17 | 10 | 0.021 | |
| ARRE_06_1164 | F | 4 | | 0.367 | X - high missingness |

| Sample ID | Sex | Modal coverage | Down-sampled modal coverage | SNP miss. freq. | Removed individual (X) |
|---|---|---|---|---|---|
| ARRE_12_1295 | M | 6 | | 0.108 | |
| ARSB_09_1050 | F | 17 | 10 | 0.026 | |
| ARSB_09_1053 | F | 7 | | 0.085 | |
| B95_51_BI | M | 3 | | 0.437 | X - high missingness |
| B96_392_SB | F | 18 | 10 | 0.027 | |
| B96_393_SB | F | 15 | 10 | 0.021 | X - kin with B96_393_SB |
| B96_394_SB | F | 20 | 10 | 0.027 | |
| B96_398_SB | F | 19 | 10 | 0.024 | |
| FMMM_RB_005_93 | F | 13 | 10 | 0.021 | |
| FMMM_RB_011_93 | M | 14 | 10 | 0.024 | |
| IMM_82_M1 | M | 18 | 10 | 0.029 | |
| MM_AB_87_005 | F | 13 | 10 | 0.024 | |
| PI_92_009_MM | M | 13 | 10 | 0.029 | |
| RA_HM_IQ_121 | M | 14 | 10 | 0.024 | X - duplicate with 94_RAHM_IQ_162 |
| RBMM01 | F | 11 | 10 | 0.025 | |
| RBMM02 | M | 14 | 10 | 0.053 | |

**Table S3**. Sample information for each bowhead whale individual, including sample ID, sex determined by genome coverage comparisons, modal coverage, final modal coverages for down-sampled samples, the frequency of SNP missingness, and notes on which individuals were removed for downstream analyses.

| **Sample ID** | **Sex** | **Modal coverage** | **Down-sampled modal coverage** | **SNP miss. freq.** | **Removed individual (X)** |
|---|---|---|---|---|---|
| 88_Pang | M | 18 | 11 | 0.019 | |
| 99_01 | F | 11 | | 0.019 | X - kin with BM_01_2009 |
| ARBMGH_2002_001 | F | 12 | 11 | 0.019 | X - kin with BM_NSA_2009_03 |
| BMDB_06_70 | F | 16 | 11 | 0.018 | |
| BMWG07_22 | F | 17 | 11 | 0.016 | |
| BMWG07_30 | M | 19 | 11 | 0.020 | |
| BM_01_2009 | F | 16 | 11 | 0.016 | |
| BM_CH_2000_01 | M | 17 | 10 | 0.024 | |
| BM_NSA_2008_02 | M | 15 | 11 | 0.020 | |
| BM_NSA_2009_02 | F | 15 | 11 | 0.014 | |
| BM_NSA_2009_03 | M | 17 | 11 | 0.022 | |
| BM_NSA_2010_01 | M | 16 | 11 | 0.021 | |
| BM_NSA_2011_01 | M | 18 | 10 | 0.022 | |
| BM_NSA_2011_03 | F | 13 | 11 | 0.014 | |
| BM_NSA_2012_02 | F | 14 | 11 | 0.015 | |
| BM_NSA_2012_03 | M | 17 | 11 | 0.020 | |
| BM_NSA_2014_01 | F | 10 | | 0.019 | |
| BM_NSA_2020_01 | M | 15 | 11 | 0.020 | |
| BM_RB_2005_001 | F | 19 | 11 | 0.016 | |
| NSA_BM_98_01 | M | 7 | | 0.092 | |
| WBF_2005_0298 | F | 14 | 11 | 0.014 | |

**Table S4**. Summary of SNP filtering steps and number of SNPs at each step for the narwhal and bowhead whale. The first part of the table includes the initial filters starting with the raw data and ending with high quality autosomal SNPs to prepare the dataset for all analyses. The second section of the table shows further filters completed for each analysis.

| Genetic variants step | Filters | Narwhal | Bowhead whale |
|---|---|---|---|
| Raw (SNPs & indels) | N/A | 7,944,818 | 15,532,800 |
| Quality filter | Removed: indels, QUAL < 30, MQ < 40, QD <4, missingness > 25%, non-biallelic sites, and scaffolds < 100 kb | 5,162,126 | 10,080,570 |
| Autosomes | Additionally removed X- and Y-linked scaffolds | 5,002,361 | 9,848,912 |

| Analysis | Additional filters | Narwhal | Bowhead whale |
|---|---|---|---|
| Population structure (pcadapt, Reich's fst) and heterozygosity (vcftools, hierfstat) | Removed sites out of Hardy-Weinberg equilibrium, minor allele frequency (0.05), and LD pruned ($r^2 > 0.8$) | 1,098,212 | 2,344,613 |
| Runs of homozygosity (plink) | Removed sites out of Hardy-Weinberg equilibrium, minor allele frequency (0.05), and LD pruned ($r^2 > 0.8$). Retained largest even scaffolds (12 for narwhal, 200 for bowhead whale) | 719,868 | 479,435 |
| Demographic history (SMC++) | Removed sites out of Hardy-Weinberg equilibrium | 4,962,230 | 9,512,046 |
| Demographic history (GONE) | Removed sites out of Hardy-Weinberg equilibrium, minor allele frequency (0.05), and retained largest even density scaffolds | 1,458,894 | 1,318,567 |
| Contemporary Ne (StrataG) | Removed sites with any missing data, minor allele frequency (0.05), and then thinned dataset to 25K SNPs | 25,000 | 25,000 |

**Table S5.** Pairwise genetic differentiation ($F_{ST}$) between each narwhal site in the eastern Canadian Arctic. AUJ = Aujuittuq; IGL = Igloolik; IKP = Ikpiarjuk; KAK = Kangiqtugaapik; KUG = Kugaaruk; MIT = Mittimatalik; NAU = Naujaat; QAU = Qausuittuq; QIK = Qikiqtarjuaq; TAL = Taloyoak

| Site | AUJ | IGL | IKP | KAK | KUG | MIT | NAU | QAU | QIK | TAL |
|---|---|---|---|---|---|---|---|---|---|---|
| **AUJ** | 0 | | | | | | | | | |
| **IGL** | 0.0056 | 0 | | | | | | | | |
| **IKP** | 0.0026 | 0.0031 | 0 | | | | | | | |
| **KAK** | 0.0016 | 0.0036 | 0.0021 | 0 | | | | | | |
| **KUG** | 0.0006 | 0.0024 | -0.0004 | 0.0006 | 0 | | | | | |
| **MIT** | 0.0020 | 0.0028 | -0.0010 | 0.0004 | -0.0013 | 0 | | | | |
| **NAU** | 0.0053 | 0.0067 | 0.0034 | 0.0031 | 0.0022 | 0.0026 | 0 | | | |
| **QAU** | 0.0007 | 0.0044 | 0.0011 | 0.0005 | 0.0004 | 0.0008 | 0.0035 | 0 | | |
| **QIK** | 0.0032 | 0.0025 | 0.0004 | 0.0010 | -0.0007 | -0.0005 | 0.0042 | 0.0017 | 0 | |
| **TAL** | 0.0002 | 0.0050 | 0.0012 | 0.0007 | -0.0007 | 0.0017 | 0.0035 | 0.00003 | 0.0021 | 0 |

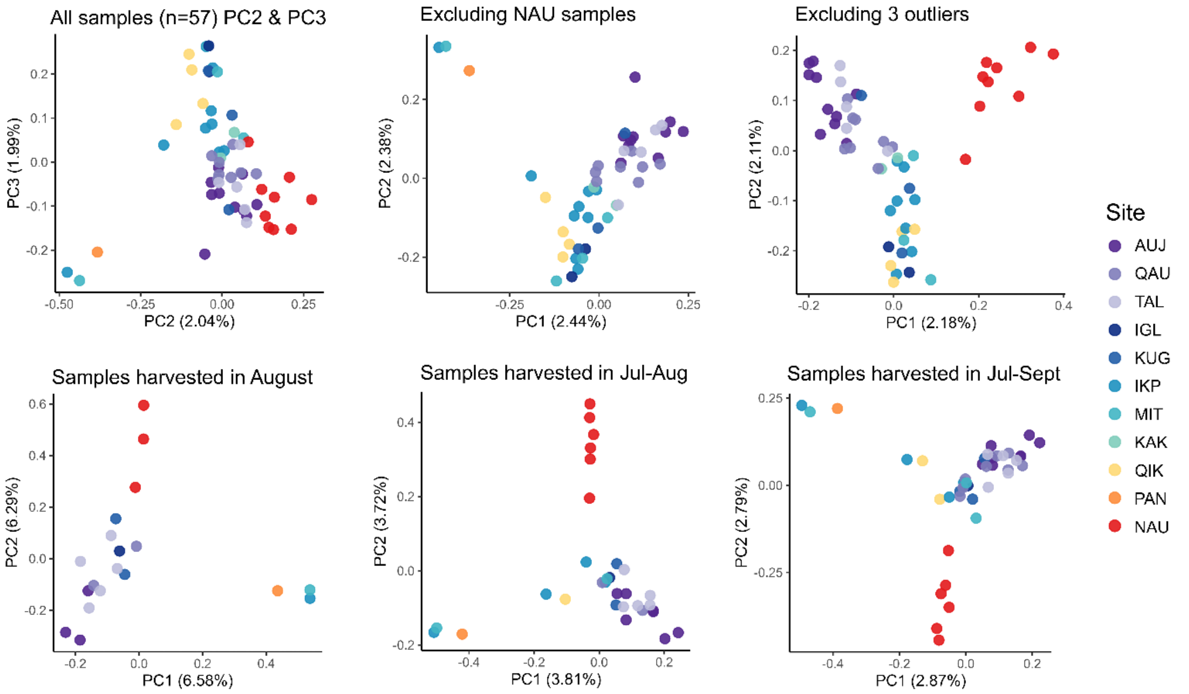


**Figure S1.** PCAs examining population structure with difference sample subsets, showing PC2 and PC3 for all samples, and PC1 and PC2 for subsets with removal of individuals from Naujaat, removal of 3 outliers, and subsets of samples based on harvest months for summer residence. Site IDs: AUJ = Aujuittuq (Grise Fiord), QAU = Qausuittuq (Resolute Bay), TAL = Taloyoak (Spence Bay), IGL = Igloolik, KUG = Kugaaruk (Pelly Bay), IKP = Ikpiarjuk (Arctic Bat), MIT = Mittimatalik (Pond Inlet), KAK = Kangiqtugaapik (Clyde River), QIK = Qikiqtarjuaq (Broughton Island), PAN = Panniqtuuq (Pangnirtung), NAU = Naujaat (Repulse Bay).

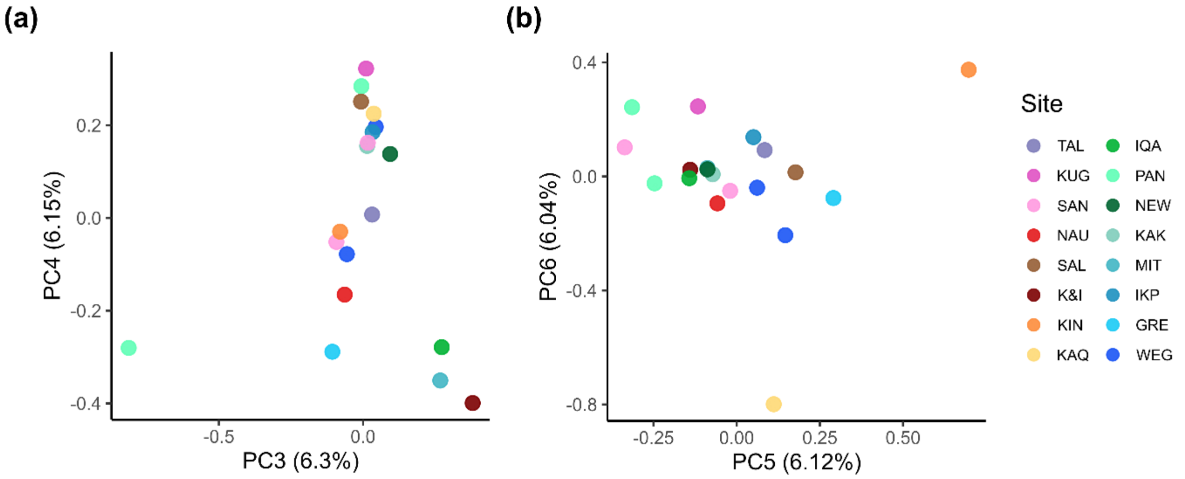


**Figure S2.** PCA results for bowhead whales (*n*=21) for (a) PC3 and PC4, and (b) PC5 and PC6. KIT = Kittigazuit (Mackenzie Delta), IKA = Ikaahuk (Sachs Harbour), TAL = Taloyoak (Spence Bay), KUG = Kugaaruk (Pelly Bay), SAN = Sanirajak (Hall Beach), NAU = Naujaat (Repulse Bay), SAL = Salliq (Coral Harbour, K&I = Kangiq&iniq (Rankin Inlet), KIN = Kinngait (Cape Dorset), KAQ = Kangiqsujuaq (Wakeham Bay), IQA = Iqaluit, PAN = Panniqtuuq (Pangnirtung), NEW = Newfoundland, KAK = Kangiqtugaapik (Clyde River), MIT = Mittimatalik (Pond Inlet), IKP = Ikpiarjuk (Arctic Bay), GRE = Greenland, WEG = West Greenland.

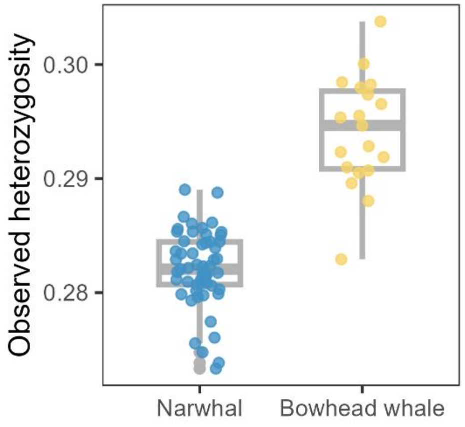


**Figure S3.** Minimal differences in observed heterozygosity in the narwhal and bowhead whale. Box plots are shown in gray, and each dot represents an individual whale.

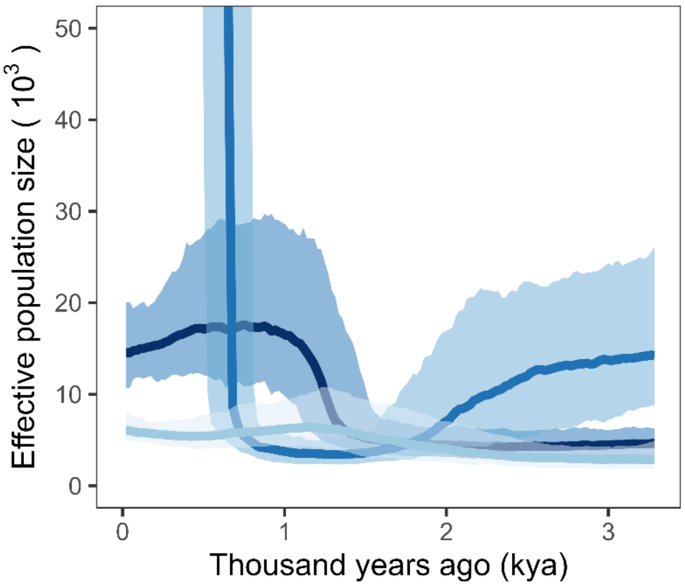


**Figure S4.** Demographic history of the last 150 generations of narwhal separated by subgroups. Dark blue = western Baffin Island, medium blue = eastern Baffin Island, and light blue = northern Hudson Bay.

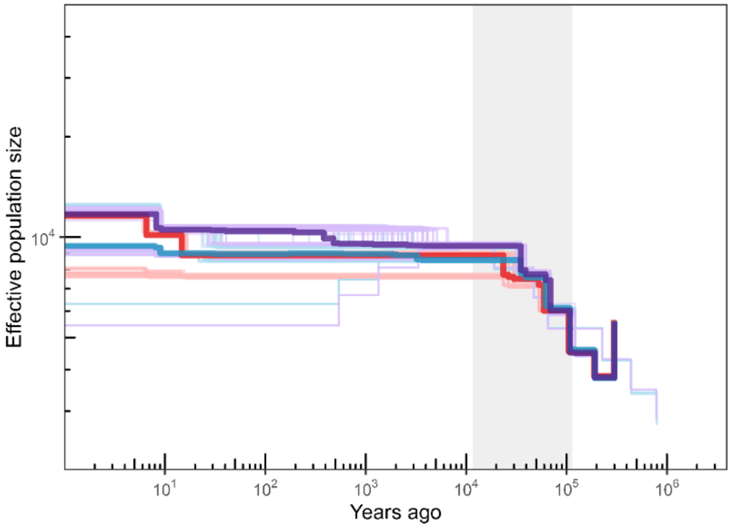


**Figure S5.** Demographic history of narwhal using SMC++ in three subgroups (purple = western Baffin Bay; blue = eastern Baffin Bay; red = northern Hudson Bay). Median runs are represented by bolded lines, and 100 iterations for each subgroup are shown in lighter tones. The last glacial period is marked by the grey bar.

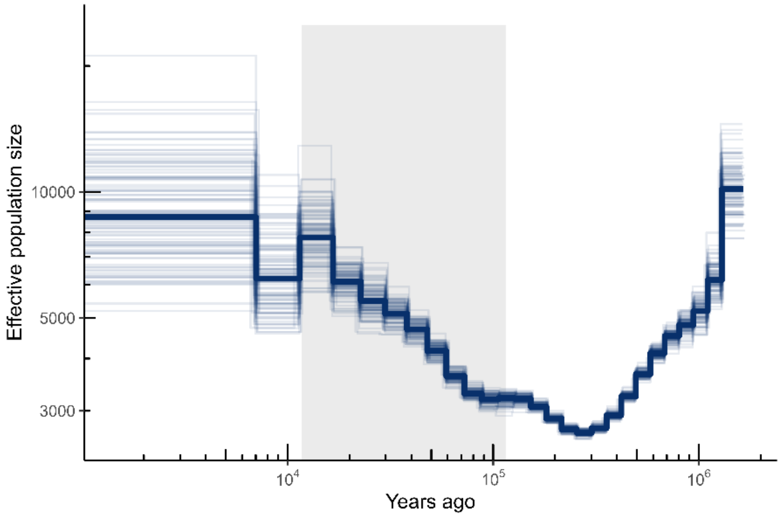


**Figure S6.** Demographic history of narwhal using PSMC. The median run is represented by the bolded line, and 100 iterations are shown in lighter tones. The last glacial period is marked by the grey bar.

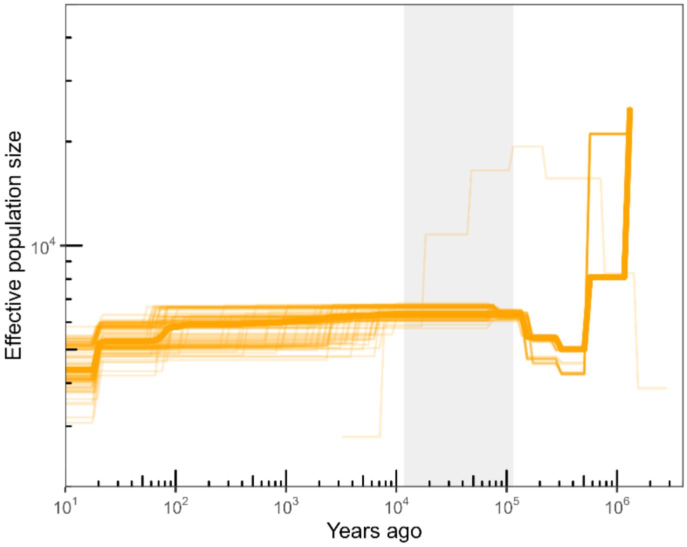


**Figure S7.** Demographic history of bowhead whale using SMC++ using all samples. Median run is represented by the bolded line, and 100 iterations are shown in lighter tones. The last glacial period is marked by the grey bar.

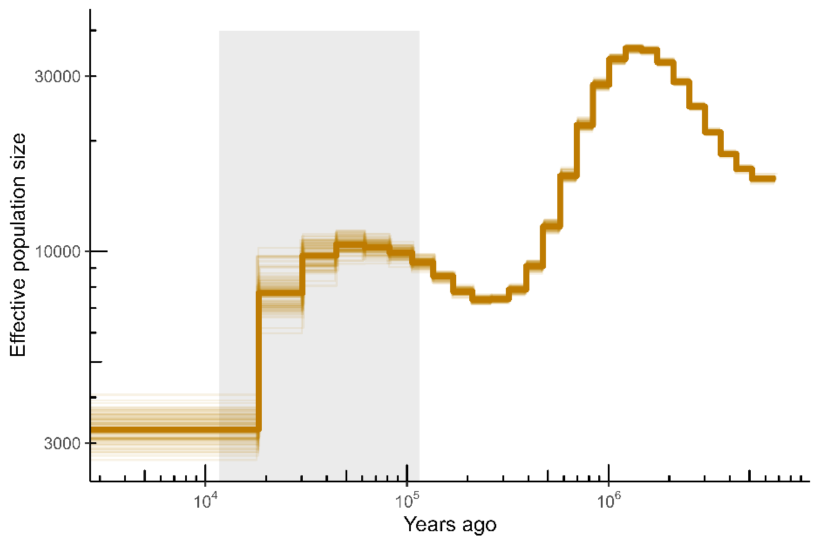


**Figure S8.** Demographic history of bowhead whale using PSMC. The median run is represented by the bolded line, and 100 iterations are shown in lighter tones. The last glacial period is marked by the grey bar.
